# Arterial endoglin does not protect against arteriovenous malformations

**DOI:** 10.1101/2020.05.23.112185

**Authors:** Esha Singh, Rachael E. Redgrave, Helen Phillips, Helen M. Arthur

## Abstract

Endoglin (ENG) forms a receptor complex with ALK1 in endothelial cells (ECs) to promote BMP9/10 signalling. Loss of function mutations in either *ENG* or *ALK1* genes lead to the inherited vascular disorder Hereditary Haemorrhagic Telangiectasia (HHT), characterised by arteriovenous malformations (AVMs). However, the vessel-specific role of ENG and ALK1 proteins in protecting against AVMs is unclear. For example, AVMs have been described to initiate in arterioles, whereas ENG is predominantly expressed in venous ECs. To investigate whether ENG has any arterial involvement in protecting against AVM formation we specifically depleted the *Eng* gene in venous and capillary endothelium whilst maintaining arterial expression, and investigated how this affected the incidence and location of AVMs in comparison with pan-endothelial *Eng* knockdown.

**Methods:** Using the mouse neonatal retinal model of angiogenesis, we first established the earliest time point at which Apj-Cre-ERT2 activity was present in venous and capillary ECs but absent from arterial ECs. We then compared the incidence of AVMs following pan-endothelial or venous/capillary-specific ENG knockout.

**Results:** Activation of Apj-Cre-ERT2 with tamoxifen from postnatal day (P) 5 ensured preservation of arterial ENG protein expression. Specific loss of ENG expression in ECs of veins and capillaries led to retinal AVMs at a similar frequency to pan-endothelial loss of ENG. AVMs occurred in the proximal as well as the distal part of the retina consistent with a defect in vascular remodelling during maturation of the vasculature.

**Conclusion:** Expression of ENG is not required in arterial ECs to protect against AVM formation.

## Background

Endoglin (ENG) is a glycosylated transmembrane protein primarily expressed on endothelial cells (ECs). Although originally thought to be a co-receptor for TGFbeta1 it is now known to have a much higher affinity for BMP9 and BMP10 ligands[1]. ENG can accumulate BMP9/10 ligand on the EC surface and supports ligand binding to the signalling receptor ALK1 (also known as ACVRL1) leading to SMAD1/5/8 phosphorylation and a range of downstream transcriptional responses [2-4]. Loss of function mutations in either *ENG* or *ALK1* genes account for the vast majority of cases of Hereditary Haemorrhagic Telangiectasia (HHT). This is an inherited vascular disorder typified by small bleeding telangiectases in the oronasal mucosa and gastrointestinal tract, as well as larger high flow arteriovenous malformations (AVMs) in lung, liver or neural tissues leading to severe morbidity [5]. HHT type I (caused by *ENG* mutations) and HHT type 2 (caused by *ALK1* mutations) are clinically very similar, except that HHT1 patients have a higher frequency of lung and brain AVMs, whilst HHT2 patients have a higher incidence of gastrointestinal bleeding [5]. The reasons for this tissue bias is unknown.

HHT1 and HHT2 patients are heterozygous for mutations in *ENG* or *ALK1* genes, respectively; although recent work has revealed that vascular lesions from HHT patients contain cells with a local second mutation in the ‘good’ allele (*ENG* or *ALK1*). This finding supports the idea that a two genetic hit mechanism is required for vascular lesion formation in HHT [6]. Strong evidence from mouse models of HHT1 and HHT2 also supports this two hit mechanism where complete endothelial loss of ENG or ALK1 protein is a pre-requisite for robust vascular lesion formation in response to angiogenic stimuli [7]. In contrast, mice that are heterozygous for mutations (*Eng*+/- or *Alk1*+/-) have low penetrance of disease [8,9].

It is well established from these studies that ENG and ALK1 proteins are required in ECs to protect the vasculature from abnormal arteriovenous shunting of blood through AVMs due to aberrant vascular remodelling. However, it is not clear whether this is primarily due to a defect in arterial or venous protective function, or both. ALK1 has been shown to be expressed primarily in adult arterial vessels [10] and loss of ALK1 in development leads to loss of expression of genes associated with arterial identity such as *Efnb2* and *Jag1* [11,12]. This suggests an arterial specific function of ALK1 signalling. Furthermore, an arteriolar origin for AVMs has been visually described in the neonatal retina following mosaic loss of ENG in ECs [13], and an arterial defect was also suggested by the finding that some arterial markers such as Ephnb2 show reduced expression in the absence of ENG [13]. In addition, there has been a long-standing debate whether AVMs in HHT are due to loss of arterial identity via disruption of the Notch pathway. There are three main lines of evidence for this idea: Notch ligands are predominantly expressed in arteries and are required to maintain arterial identity, loss of Notch activity leads to AVMs, and Notch signalling synergises with ALK1 signalling [14-17]. Therefore, to address whether there is arterial involvement in the formation of AVMs in HHT, we depleted ENG specifically in venous and capillary ECs, leaving arterial ENG expression intact. We then analysed AVM formation using the well-established neonatal mouse model of retinal angiogenesis.

## Methods

The care and use of animals were in accordance with the UK Government Animals (Scientific Procedures) Act 1986 and were approved by Newcastle University’s Animal Welfare and Ethical Review Body. *Eng*^*fl/fl*^ mice[18] were intercrossed with *Cdh5-Cre-ERT2* [19] or *Apj-Cre-ERT2* [20] lines to permit pan-endothelial or venous/capillary specific targeting of the ENG floxed allele, respectively. To map Apj-Cre-ERT2 activation we used the *Rosa26-mTmG* Cre reporter line [21] and treated neonates with a single subcutaneous injection of tamoxifen (Sigma, T5648) dissolved in peanut oil: ethanol (10:1) at postnatal day (P)2 (0.5mg), P4 (0.5mg) or P5 (0.6mg). To generate efficient recombination of the *Eng* floxed allele, pups were given 0.6mg tamoxifen on consecutive days at P5 and P6. Retinas were then harvested at different time points as indicated in the main text.

Mouse neonatal retina analysis was performed using a similar approach to our earlier work [22] and described in detail elsewhere [23]. Essentially, following enucleation of the eye and dissection, the retina was imaged directly to record RFP and GFP staining under an M2 Axio-imager microscope using an Axiocam digital camera. Subsequently retinas were subject to methanol fixation and staining with isolectin B4-Alexa488 (Life Technologies) and antibodies against ENG (anti-CD105 from eBioscience), and smooth muscle actin (anti-aSMA-Cy3 from Sigma Aldrich). Secondary anti-rat antibody conjugated to Alexa568 or Alexa 647 (Life Technologies) was used to detect anti-CD105. Images were captured using an M2 axio-imager microscope (Zeiss) and staining intensity analysis was performed using Zen software. For depth coding, z-stack images of the primary and secondary plexus were colour coded according to the distance from the upper surface of the retina using Zen software. For all experiments either one or both retinas from each pup were stained and data was analysed using Graphpad prism software.

## Results

Our first goal was to establish a mouse model of angiogenesis in which arterial endoglin could be maintained whilst depleting ENG in venous and capillary ECs. The *Apj-Cre-ERT2* mouse line shows venous/capillary endothelial specific expression and has been used successfully to track venous ECs in coronary vessel development [20]. However, to establish when Cre-ERT2 expression from the Apj promoter becomes venous endothelial-specific during development of the neonatal vascular retinal plexus, we used the *Rosa26-mTmG Cre* reporter mouse [21]. CRE-mediated recombination causes the mTmG reporter expression to switch from tomato red fluorescent protein (RFP) to green fluorescent protein (GFP) allowing localisation and timing of CRE activity to be determined. Treatment of *Apj-Cre-ERT2*;*R26-mTmG* neonates with tamoxifen at different postnatal ages (up to and including P5) and retinal tissue analysis at P8 showed that tamoxifen treatment at P5 led to arterial ECs retaining RFP expression whilst venous and capillary ECs became GFP-positive due to CRE-mediated recombination of the *R26-mTmG* reporter(Supplementary Fig. S1). In contrast, neonates that had been injected with tamoxifen at earlier time points (P2 or P4) showed significant GFP expression in the arteries reflecting arterial Apj-Cre activity. Thus, P5 was the earliest time point at which Apj-Cre-ERT2 activation becomes specific to non-arterial ECs in the developing retina. This is in agreement with the previously reported age of P5 when *Apj* transcripts are lost from neonatal retinal arteries [24]. This timing is likely to be because it takes 5 days of postnatal retinal vascular development for the arterial endothelium to become phenotypically established, combined with the fact that pre-existing venous ECs can contribute to the arterial endothelium during retinal development via a process of EC migration against blood flow [25,26].

We have previously shown that pan-endothelial knockout of ENG (Eng-iKO^e^) using *Eng*^*fl/fl*^;*Cdh5-Cre-ERT2* neonates led to AVMs affecting 70% of Eng-iKO^e^ retinas [22]. However, initiation of Eng depletion was at P2, which is substantially prior to the establishment of differentiated arteries and veins in the developing retina. Therefore, to establish the phenotype of pan-endothelial Eng knockout at the later time points required for arterial specific Apj-Cre activity, we first determined whether delivery of tamoxifen at P5 and P6 to *Engfl/fl;Cdh5-Cre-ERT2* neonates led to retinal AVMs. Harvesting the tissue at P8 revealed that ENG protein was only partially depleted (not shown), likely because the time between Cre-ERT2 mediated recombination of the *Eng* floxed gene and tissue harvest was insufficient to allow complete turnover of pre-existing ENG protein. We therefore repeated the experiment leaving additional time for ENG protein loss, this time harvesting retinas at P11. In this scenario tamoxifen activation of *Eng*^*fl/fl*^;*Cdh5-Cre-ERT2* led to efficient ENG protein knockdown and the proportion of Eng-iKO^e^ retinas with AVMs was 67% (Fig 1c), similar to our previous findings [22]. To compare this outcome with the effect of retaining arterial ENG expression, whilst knocking out ENG in venous and capillary ECs, *Eng*^*fl/fl*^;*Apj-Cre-ERT2* neonates were treated with tamoxifen at P5 and P6 and tissue was harvested at P11 to generate venous and capillary specific Eng knockout (Eng-iKO^v^) mice. Immunostaining of Eng-iKO^v^ retinas confirms ENG protein depletion in capillaries and veins, whilst ENG expression is retained in arteries (Fig. 1e). Over 90% of Eng-iKO^v^ retinas show AVM formation (Fig. 1b,c), pointing to the absence of a role for arterial ENG in protecting against AVM formation. Littermate *Eng*^*fl/fl*^ controls treated with tamoxifen retain ENG expression in all vessels and have no AVMs (Fig.1a,d).

**Fig. 1.**
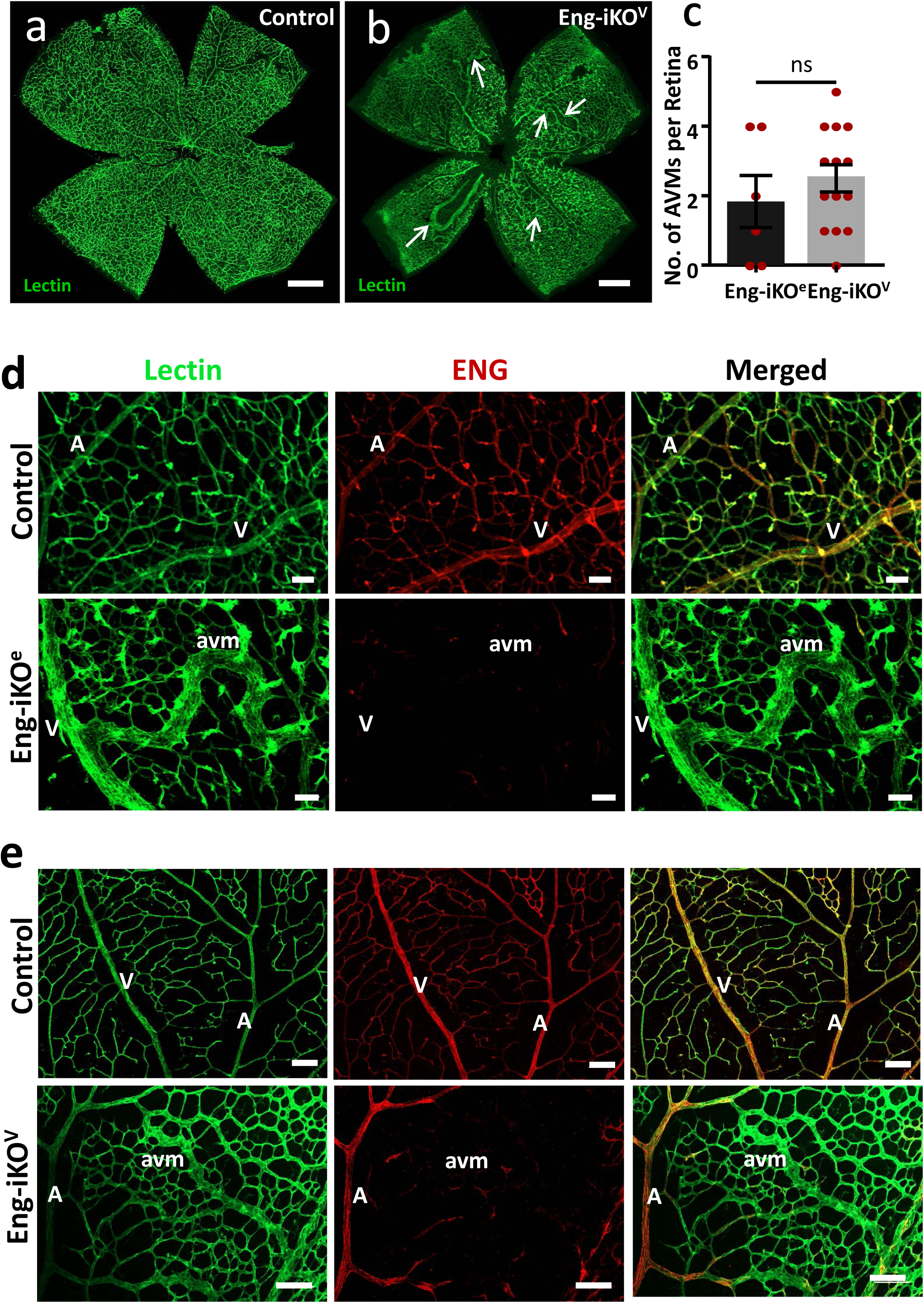
Pan-endothelial or venous-capillary specific loss of ENG expression leads to similar incidence and frequency of AVMs. **a:** Control retina harvested at P11 show the normal radial distribution of arteries and veins, with intervening capillaries. **b:** Loss of ENG in venous and capillary ECs in Eng-iKO^v^ retinas leads to multiple AVMs (arrows). Scale bar = 500μm. **c:** Incidence of AVMs at P11 is similar between Eng-iKO^e^ (n=6) and Eng-iKO^v^ (n=16) retinas and the mean number of AVMs per retina is 1.8 in Eng-iKO^e^ neonates and 2.5 in Eng-iKO^v^ retinas. The frequency of P11 retinas that have AVMs is 67% (4/6) in Eng-iKO^e^ neonates and 94% (15/16) in Eng-iKO^v^ neonates. Data were analysed by unpaired t test. **d:** Eng-iKO^e^ retinas were stained with isolectin and anti-ENG antibody. Pan-endothelial loss of ENG in Eng-iKO^e^ retinas leads to AVM formation. **e:** Eng-iKO^v^ retinas retain arterial ENG protein, but develop AVMs in a similar way to Eng-iKO^e^ retinas. Scale bar =50μm. Abbreviations: A, artery; V, vein; avm, arteriovenous malformation

There were no detectable differences between the vascular phenotypes following venous-specific or pan-endothelial loss of ENG and venous enlargement was pronounced “downstream” of the AVMs, likely in response to increased blood flow (Supplementary Fig. S2a). The most distal portion of the arteries, where they branch to form capillaries close to the periphery, shows evidence of mTmG recombination when Cre is activated at P5 (Supplementary Fig. S1a,b,c). Therefore, it was important to establish whether AVMs were preferentially located at the most distal part of the artery, where ENG expression was lost, or in the mid- and proximal regions where ENG protein was retained. Measuring the position of each AVM with respect to the arterial distance from the centre of the retina, and comparing this with the arterial length where ENG protein expression was retained, we found that AVMs were primarily located in the mid and proximal regions of the retina, and adjacent to ENG expressing arteries (Fig. 2a,b,c). To investigate whether normal levels of ENG protein expression were maintained on the arterial side of these AVMs we performed immunofluorescence staining intensity analysis. This confirmed that ENG protein levels were unaffected in arteries of Eng-iKO^v^ retinas compared with controls, whilst ENG was significantly reduced in capillaries, veins and AVMs (Fig. 2d). AVMs were muscularised (Fig. 2a) likely in response to increased blood flow, similar to our previous observations [22].

**Fig. 2.**
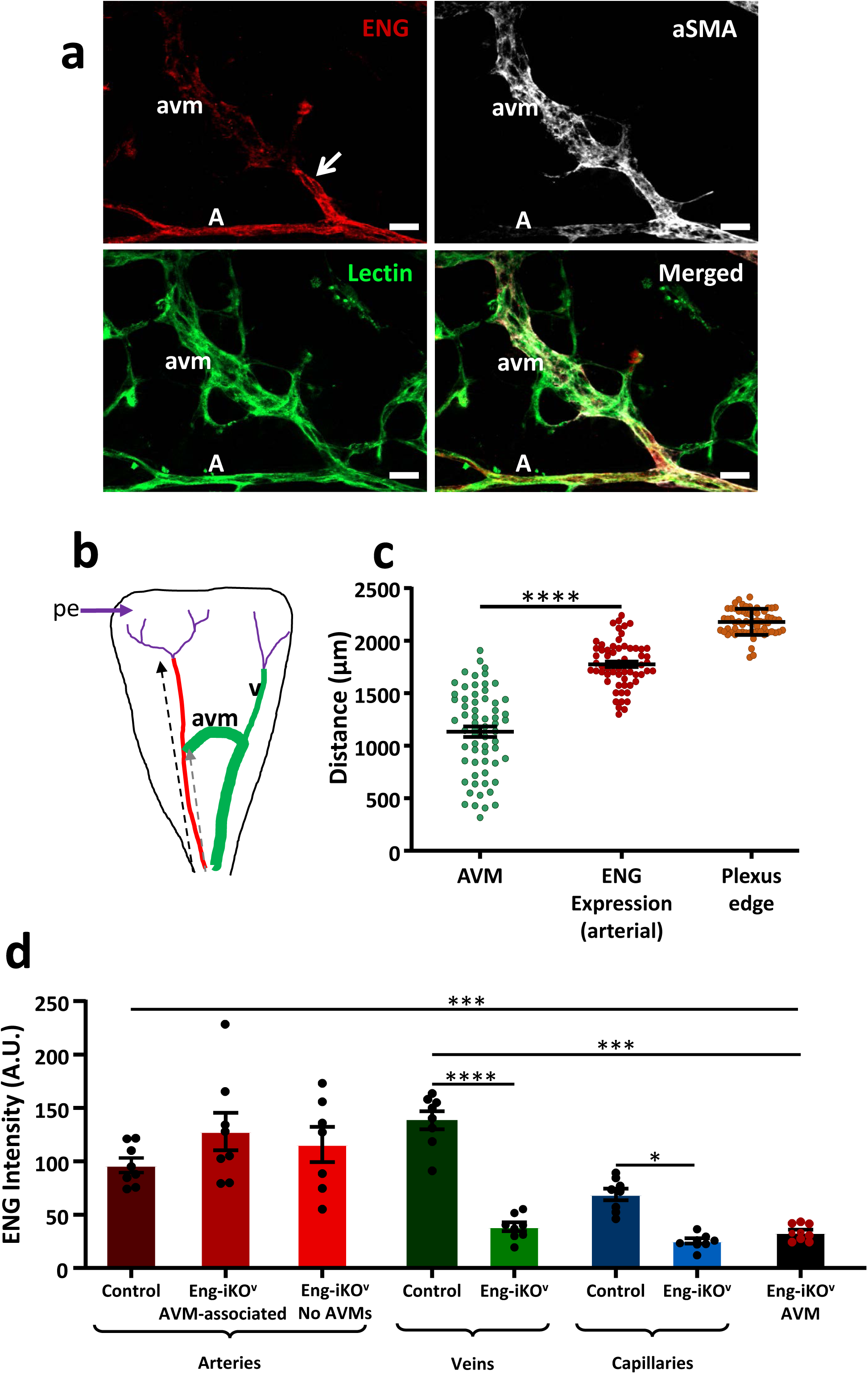
ENG expression is maintained on the arterial side of AVMs in Eng-iKO^v^ retinas. **a**: Vascular endothelial cells in P11 Eng-iKO^v^ retinas were identified using isolectin staining. These retinas show ENG-positive arteries, whilst AVMs and capillaries show reduced ENG staining. AVMs and arteries also have vascular smooth muscle coverage (aSMA). A total of 42 AVMs were analysed and in all cases ENG protein was maintained on the arterial side of the malformation (arrow). Scale bar=20μm. **b**: Cartoon to illustrate the extent of ENG expression (red) in arteries measured from the centre of the retina to the site at which arterial ENG expression is lost (black dashed arrow), which is close to the vascular plexus edge. Arterial ENG expression extends beyond the point at which AVMs form (grey dashed arrow). Veins (v) and AVMs are shown in green, and capillaries in purple, with position of plexus edge (pe) marked (purple arrow). **c:** The distance from the centre of the retina to the arterial site of each AVM was measured. This was compared with the arterial distance at which ENG expression is lost, and shows that AVMs originate from ENG expressing arteries and AVMs were primarily located in the mid-plexus region. The distance to the plexus edge is shown to provide context for the analysis. The arterial origin of AVMs was significantly closer to the centre of the retina than the site at which arterial ENG protein was lost (n=16/group). Data were analysed by paired t-test. **** p<0.0001. **d:** ENG immunofluorescent staining intensity was measured in different vessel types in Eng-iKO^v^ retinas at P11 (n=9). ENG protein is retained at control levels in Eng-iKO^v^ arteries whether AVMs were present or not, but ENG protein is significantly reduced in veins, capillaries and AVMs. Data were analysed by one way ANOVA with Bonferroni correction for multiple comparisons. *p<0.05; *** p<0.001. Abbreviations: A, artery; V, vein; avm, arteriovenous malformation

A potential complication at P11 is that branches of the upper vascular plexus have begun to migrate down into the retina to form the deeper vessel plexus [27,28]. We therefore used depth coding to locate AVMs and confirmed they were in the primary plexus (Fig 3). Also, as the primary plexus had already reached the retinal periphery by P8, prior to full ENG protein knockdown, and AVMs were mainly found in the mid-plexus region, this suggests that AVMs form in the remodelling vasculature of the Eng-iKO^v^ retinas and are not dependent on active angiogenic processes at the leading edge of the developing vascular plexus.

**Fig. 3.**
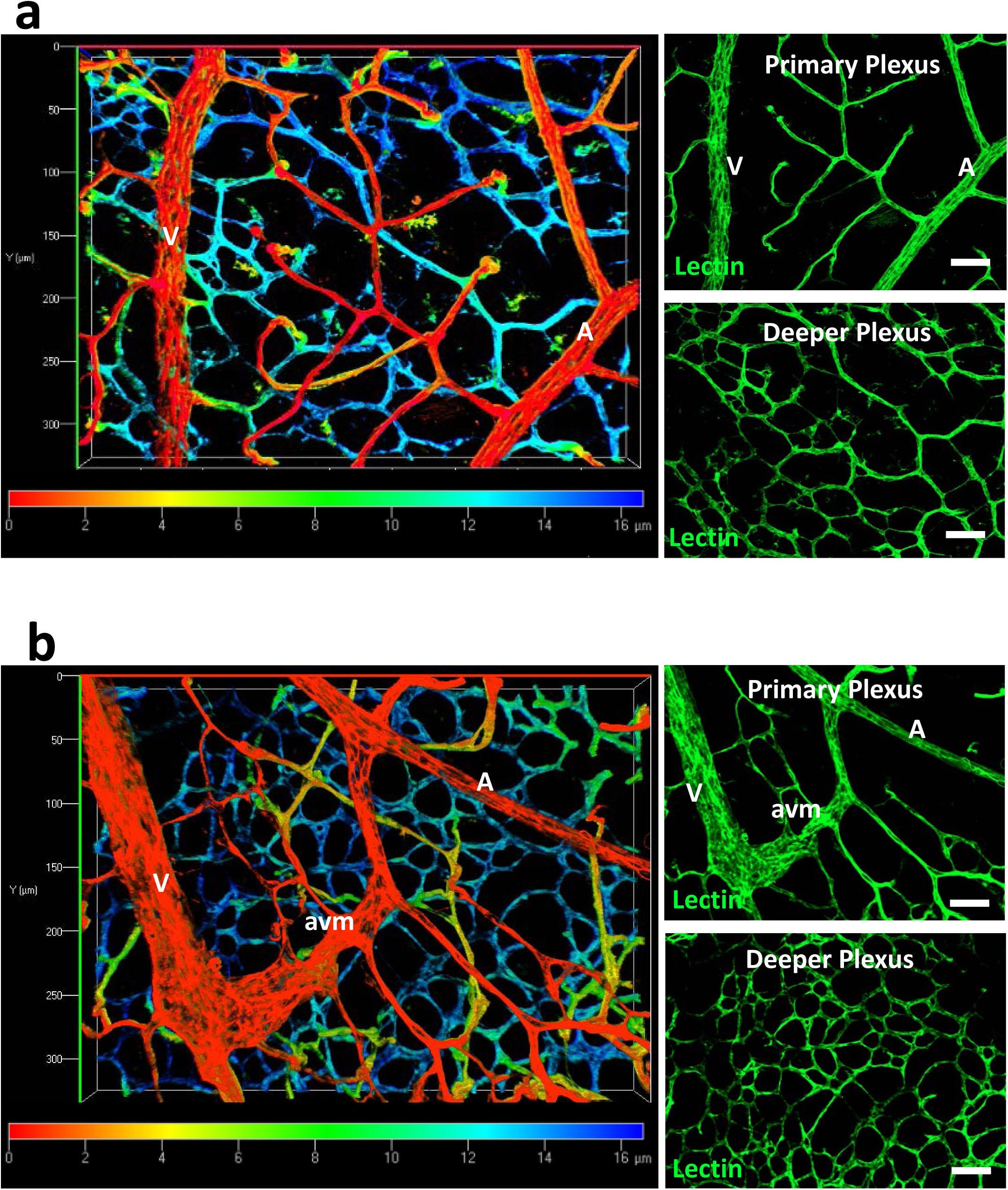
AVMs in Eng-iKO^v^ retinas are in the primary vascular plexus. **a:** Depth coding images of isolectin stained control retinas at P11 (n=20) allow discrimination of the primary and deeper plexus. Colour spectrum scale denotes relative depth of focus with red corresponding to vessels closest to the retinal surface (primary plexus) and blue corresponding to the vessels lying deeper in the retinal tissue. Separate Z stack images of the primary and deeper plexus are shown for comparison. **b:** Depth coding analysis confirms that AVMs are located in the primary plexus of P11 retinas (n=16) harvested from Eng-iKO^v^ neonates (following tamoxifen treatment at P5 and P6). Scale bar: 50μm. Abbreviations: A, artery; V, vein; avm, arteriovenous malformation.

## Discussion

There is no difference in incidence or location of AVMs in venous-specific and pan-endothelial Eng-iKO retinas suggesting that loss of arterial ENG is not involved in driving the formation of AVMs. Instead, AVMs result from ENG defects in venous and/or capillary ECs. Furthermore, AVMs can form in the retinal primary plexus after the main neo-angiogenesis events are complete and arteries have become muscularised. This finding suggests that AVMs are not restricted to areas of neo-angiogenesis, but can also occur in regions of vessel remodelling and maturation. We have recently reported a similar scenario where aberrant remodelling of mature peripheral vessels of the adult mouse leads to major AVMs following pan-endothelial ENG depletion [29]. Further work is required to determine whether loss of ENG in mature vessels leads to AVMs via reduced EC migration against flow, a process previously shown to underpin AVM formation in actively angiogenic vessels [13], or whether an abnormal increase in EC proliferation is the primary cause of AVMs in pre-established blood vessels.

Our work also helps to address a long-standing controversy regarding whether AVMs form because of loss of arterial identity. This idea has come from the fact that Notch signalling maintains arterial identity and loss of function Notch mutations lead to AVMs [16,17]. Furthermore, Notch and ALK1 signalling activities synergise, in that SMAD1/5/8 activity promotes transcription of Notch gene targets, such as those required for arterial identity. It has therefore been proposed that HHT may result from a loss of arterial identity due to reduced Notch activity. However, although crosstalk between Notch and Smad1/5/8 signalling is critical for the stalk cell phenotype [14] it does not appear that arteries play a direct role in AVM formation in HHT. We show here that even when arteries are unaffected, AVMs develop in the same way and at a similar frequency to pan-endothelial depletion of ENG. This finding is in agreement with a venous and angiogenesis role for ENG, rather than an arterial role, and is supported by the fact that ENG expression is stronger in veins and proliferating capillaries compared with arteries [22,13]. We find the increased venous expression of ENG is maintained at P11, although ENG expression in the established capillaries of P11 retinas is now reduced below that of arteries (Supplementary Fig. 2b). Furthermore, we recently showed that the AVM phenotype in Eng-iKO^e^ mouse retinas persists despite overexpression of human (h) ENG in arteries, and it was not until hENG was also overexpressed in veins and capillaries that the AVM phenotype was rescued [13]. Our findings also align with recent work in zebrafish showing that Alk1 is not required for arterial identity, and AVM formation in *alk1* mutants does not result from loss of Notch activity [30]. As the molecular role of ENG is to promote ALK1 signalling by facilitating ligand binding it is likely that ALK1, like ENG, will also not have an arterial role in protecting against AVMs in mammals, although supporting evidence currently awaits further studies. It is interesting to consider in this context our original observation that ENG expression is decreased in *Alk1-iKO*^*e*^ retinas [12]. This raises the question of whether the reverse is true such that ALK1 expression is reduced in *Eng-iKO*^*e*^ retinas. In fact we observed that ALK1 expression appears to increase in the high flow AVMs of Eng-iKO^e^ retinas using our previous knockout protocol [22] (Supp Fig 3). These findings are entirely consistent with previous studies showing that ENG is downstream of BMP9 signalling and is therefore be reduced in the absence of ALK1 [31,12], whilst in contrast, ALK1 expression has not been reported to be regulated by BMP9, but is upregulated by sheer stress [32,33], which is anticipated to be increased due to the high blood flow through AVMs. Although endothelial loss of ENG or ALK1 lead to AVMs in preclinical models, the phenotype of ALK1-iKO^e^ mutants is more severe than that of Eng-iKO^e^ mutants [7]. This is likely because loss of ALK1, the signalling receptor, leads to elimination of this signalling pathway, whilst loss of ENG, the co-receptor, reduces the availability of ligand to the signalling receptor [3], thereby dampening rather than completely obliterating signal. In HHT patients however, the differences between HHT1 (*ENG*) and HHT2 (*ALK1*) clinical phenotypes are more complex and await a better understanding of the relative contribution of second hits and additional factors such as inflammation to fully unravel the etiology of this disease.

Finally, we show for the first time that AVMs can form in the murine retinal blood vessels during the second week of life, after the primary plexus is established, which considerably extends the previous time frame available for these studies, opening up further options for therapeutic rescue experiments.

## Acknowledgements

This work was funded by the British Heart Foundation (RG/12/2/29416, PG/18/25/33587, PG/18/32/33760) and the Cookson Trust. We are grateful to Kristy Red-Horse for sharing the Apj-Cre-ERT2 mouse line.

## Declarations

The authors declare no conflict of interest; primary data is available from the corresponding author upon reasonable request.

## Notes

### Competing Interest Statement

The authors have declared no competing interest.

