## Supplementary Figures for "Arterial endoglin does not protect against arteriovenous malformations"

#### Figure Legends

**Supplementary Fig. 1 Apj-Cre-ERT2 activity is lost from the arteries at P5**.

**a:** *Apj-Cre-ERT2;R26-mTmG* mice were injected with tamoxifen to activate Cre at different postnatal ages and retinal tissue was harvested at P8 for analysis. Retinas from neonates injected with tamoxifen at P2 (n=2) or P4 (n=2) showed significant Cre mediated recombination to generate GFP expression (green) throughout the vasculature including arteries (arrows). In contrast, tamoxifen treatment at P5 resulted in no Cre activity in ECs of the main arteries, which retained RFP expression (red) whilst venous and capillary ECs became GFP-positive (n=5). The transition point at which distal arterial GFP expression is present is indicated (arrowheads) and corresponds to the site at which the arteries branch to become capillaries close to the plexus periphery. Scale bar: 200µm. Abbreviations: v, vein; a, artery.

**b:** The distance from the centre of the retina to the point on the artery at which GFP is first seen is shown for retinas from *Apj-Cre-ERT2;R26-mTmG* injected with tamoxifen at P5 and P6 and harvested at P8 (n=7). This transition is very close to the full artery length prior to its branching to form capillary branches at the retina periphery.

**c:** The same measurements as B were made for *Apj-Cre-ERT2;R26-mTmG* retinas (n=2) from neonates injected with tamoxifen at P5 and P6 and tissues harvested 3 days later at P11.

Abbreviations: A, artery; V, vein.

**Supplementary Fig. 2**

**a:** Eng-iKOe and *Eng-iKO*$^{v}$ retinas harvested at P11 show a very similar phenotype, and have significantly enlarged AVMs. Venous diameter in both mutants is also significantly enlarged in regions where blood is shunted via an AVM (veins post AVM), compared to veins fed by normal capillary drainage. Data were analysed by two way ANOVA. There was no significant difference between genotypes, but colour coded # indicates significantly ($p<0.0001$) enlarged vessels with respect to control veins within each genotype.

**b:** Control (*Eng*$^{fl/fl}$ ) retinas at P11 show highest ENG protein expression in veins compared with arteries and established capillaries. Data were analysed by one way ANOVA with Bonferroni correction for multiple comparisons. *$p<0.05$; ** $p<0.01$; ****$p<0.0001$.

**Supplementary Fig. 3**

*Engfl/fl* (control) and *Cdh5-Cre-ERT2;Engfl/fl* (Eng-iKOe) neonates were injected with tamoxifen at P2 and P4 to activate Cre and retinal tissue was harvested at P8 for analysis, similar to our previous work [22]. Control retinas show ALK1 expression in arteries, veins and capillaries as previously shown [12]. Eng-iKOe retinas show strong ALK1 expression in arteries, veins and AVMs. ALK1 expression is high in AVMs, potentially in response to increased blood flow. Scale bar: 100μm. Abbreviations: v, vein; a, artery; avm, arteriovenous malformation.

### Supplementary Fig. S1

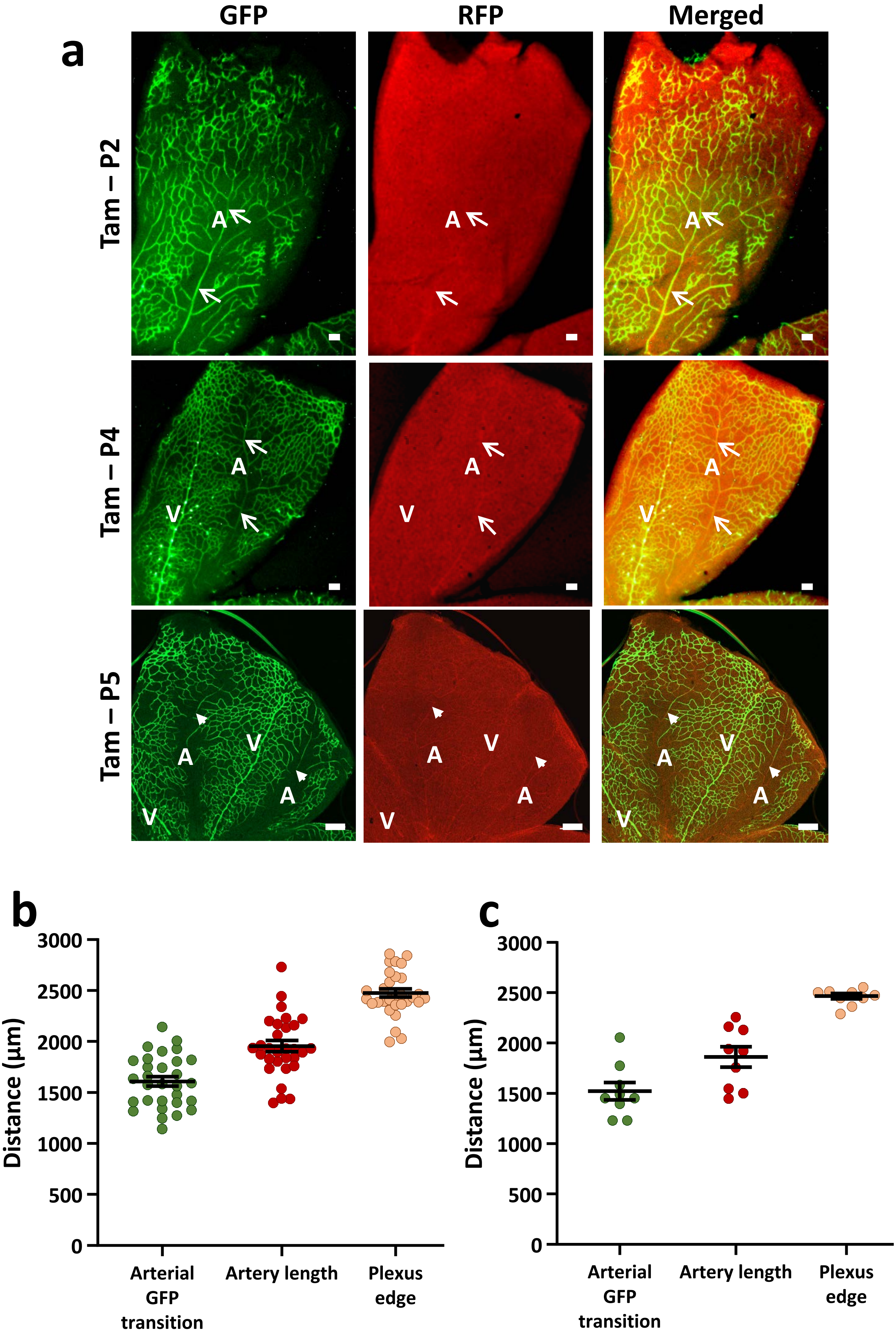

### Supplementary Fig. S2

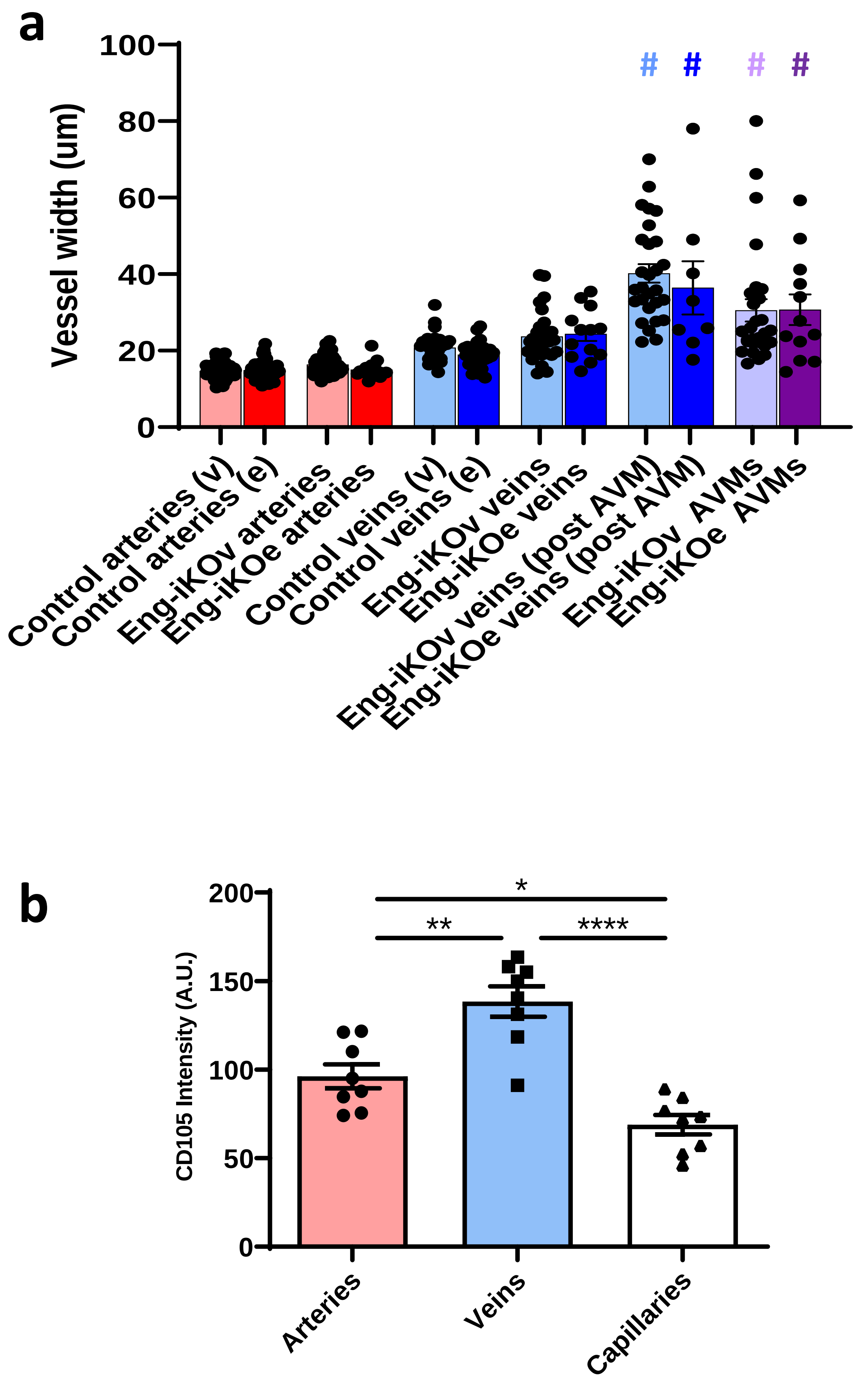

### Supplementary Fig. S3

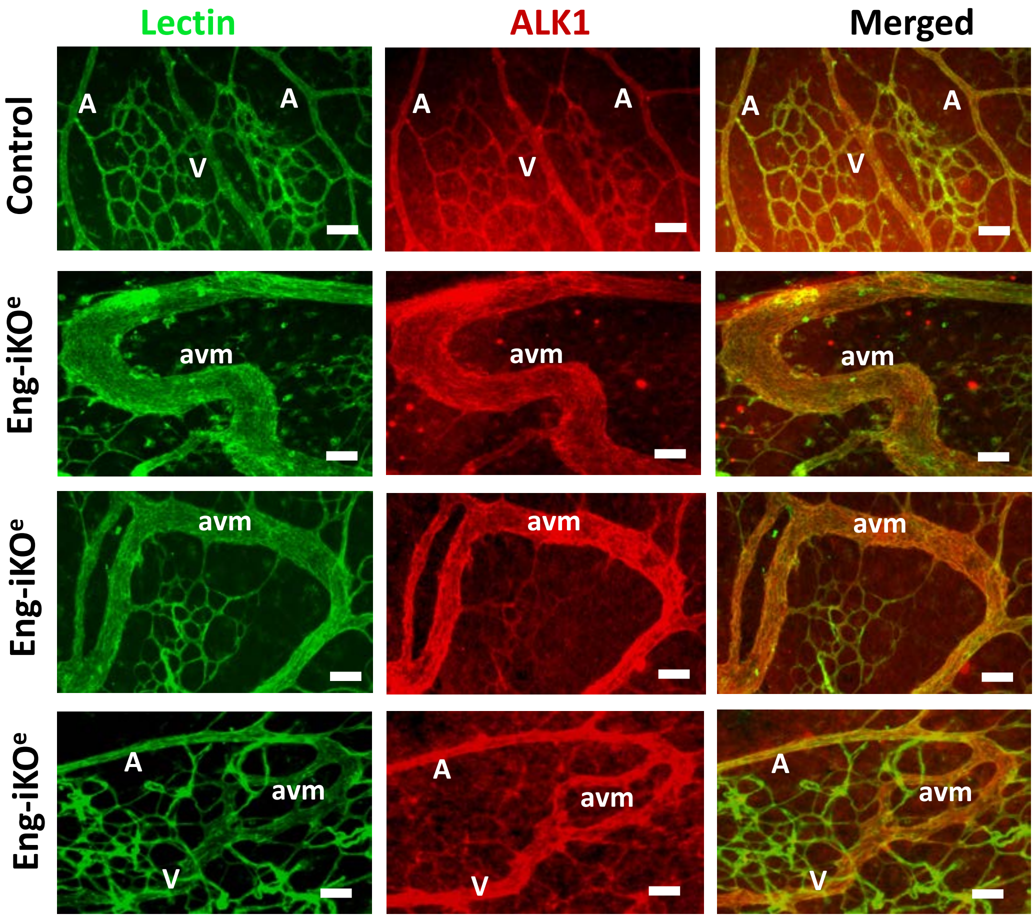
